# Engram reactivation during memory retrieval predicts long-term memory performance in aged mice

**DOI:** 10.1101/2020.01.12.903088

**Authors:** Kubra Gulmez Karaca, David V.C. Brito, Janina Kupke, Benjamin Zeuch, Ana M.M. Oliveira

## Abstract

Age-related cognitive decline preferentially targets long-lasting episodic memories that require intact hippocampal function. Memory traces (or engrams) are believed to be encoded within the neurons activated during learning (neuronal ensembles), and recalled by reactivation of the same population. However, whether engram reactivation dictates memory performance late in life is not known. Here, we labelled neuronal ensembles formed during object location recognition learning in the dentate gyrus, and analyzed the reactivation of this population by long-term memory recall in young adult, cognitively impaired-and unimpaired-aged mice. We found that reactivation of memory-encoding neuronal ensembles at long-term memory recall was disrupted in impaired but not unimpaired-aged mice. Furthermore, we showed that the memory performance in the aged population correlated with the degree of engram reactivation at long-term memory recall. Overall, our data implicates recall-induced engram reactivation as a prediction factor of memory performance in aging. Moreover, our findings suggest impairments in neuronal ensemble stabilization and/or reactivation as an underlying mechanism in age-dependent cognitive decline.

## 1. Introduction

Age-related cognitive decline refers to the gradual decrease in cognitive performance throughout the aging process, mostly affecting long-term storage of episodic and spatial memories that depend on hippocampal function (Burke and Barnes, 2006). The dentate gyrus (DG) is the hippocampal subregion most sensitive to the effects of advanced age (Small et al., 2011, 2004). The DG undergoes anatomical and physiological changes as well as alterations in the transcriptomic profile that are thought to underlie the aging-associated dysfunction (Burke and Barnes, 2010; Ianov et al., 2016). It has recently been shown that subsets of DG neurons activated during learning (i.e., neuronal ensembles) belong to the memory engram, and that the reactivation of this neuronal population at memory recall is necessary and sufficient to evoke memory retrieval in young adult mice (Josselyn et al., 2015; Tonegawa et al., 2015). Moreover, dysfunctions in neuronal ensemble reactivation and memory retrieval mechanisms have been proposed as an underlying cause of dementia in pathological conditions, such as Alzheimer’s disease (Perusini et al., 2017; Poll et al., 2020; Roy et al., 2016), in which aging presents a major risk factor. However, whether memory impairments observed late in life are associated with disruptions in neuronal ensemble reactivation is not known.

Here, we hypothesized that differences in DG neuronal ensemble dynamics underlie the inter-individual variability in long-term memory (LTM) performance in the aged population. We used tagging tools to label and characterize the formation and reactivation of neuronal ensembles associated with long-term spatial recognition memory, a form of memory particularly susceptible to decline with age. We showed that impairments in long-term spatial memory during aging are associated with impaired reactivation of memory-encoding neuronal ensembles at LTM recall. Moreover, we found that the degree of engram reactivation in aged mice correlated with long-term spatial memory performance. Overall, our findings identified novel cellular correlates of age-related memory decline.

## 2. Materials and Methods

### 2.1. Animals

8-10 weeks or 18-20 months old male C57BL/6JRj mice (Janvier Labs, Germany) were individually housed on a 12 h light/dark cycle and *ad libitum* access to food and water (after stereotaxic surgery regular food was replaced by doxycycline-containing diet (40mg/kg, BioServ, Flemington, NJ, USA)). Experiments were carried out during the light phase. All of the animal procedures were approved and performed according to the European Community Council Directive 86/609/EEC.

### 2.2. Recombinant adeno associated viruses (rAAVs)

rAAVs were produced and purified as described previously (Gulmez Karaca et al., 2020; Zhang et al., 2007). The RAM-HA-GFP viral construct was generated by insertion of a HA-tagged GFP expression cassette into the multiple cloning site (MCS) of rAAV-RAM (rAAV-RAM-d2TTA::TRE-MCS-WPRE-pA) that was a kind gift from Yingxi Lin (Addgene plasmid # 63931; http://n2t.net/addgene:6393; RRID:Addgene_63931)) (Gulmez Karaca et al., 2020; Sørensen et al., 2016).

### 2.3. Stereotaxic surgery

250 nl of rAAV-RAM viral solution was injected into the dentate gyrus (DG) of the hippocampus at the following coordinates relative to Bregma:-2.0 mm anteroposterior, ± 1.3 mm medio-lateral, - 2.4 mm dorsoventral at the rate of 80-100 nl / min. The needle was left in place for 10 min before and after each injection to allow DG-specific diffusion of the fluid. Only the mice with viral spread throughout the DG of at least 60 µm range in the AP axis were considered for behavior and further analysis. Throughout the study, a total of 10 mice were rejected from the analysis due to insufficient viral spread (5 mice) and/or lack of exploration during behavioral training or testing (5 mice).

### 2.4. Behavior paradigms

Three weeks after stereotaxic surgery and following 5 days of handling, the spatial object recognition (SOR) test was performed. SOR training involved four sessions; each lasted 6 min and was separated by 3 min intervals in the home-cage. In the first session, mice explored an empty black, square open field (50 cm × 50 cm × 50 cm) with a visual cue placed on one wall. This session was also used for the open field test as described previously (Brito et al., 2020b, 2020a; Gulmez Karaca et al., 2018). The Smart Video Tracking Software (Panlab, Harvard Apparatus) was used to score the time spent in the central zone (32% of the arena), number of center entries and total distance travelled in the open field test. In the next three sessions of the SOR training, mice freely explored two diagonally located distinct objects (a glass bottle and a metal tower) in the arena. After the training, mice were placed in the home-cage and were undisturbed until the memory test. During the memory test (24 h post-training), one of the two objects was displaced in the arena (while the other one remained in the original position), which the mice explored freely for 6 min. Object exploration was defined as the animal sniffing the object or pointing its nose to the object within 1-cm distance. When the animal leaned on the object but did not direct the nose towards it, object exploration was not considered. SOR memory performance was assessed by the formula (T_displaced_ /(T_nondisplaced_ + T_displaced_) × 100) where T represents time exploring an object. Assigning the aged mice into cognitively impaired and unimpaired groups was performed based on the mean SOR memory performance of the young group. Aged mice displaying memory performance lower than one standard deviation from the mean of young mice were designated as cognitively impaired.

### 2.5. Immunohistochemistry

2 h after the start of the SOR test, mice were perfused intracardially with 4% paraformaldehyde (PFA) (Sigma, Munich, Germany) and free-floating brain slices (at a thickness of 20 µm) were immunostained as previously described (Gulmez Karaca et al., 2018; Oliveira et al., 2012). Primary antibodies were used at the following concentrations: anti-HA tag (Covance MMS-101R (1:1000)), anti-Fos (Cell Signaling #2250, 1:1000)).

### 2.6. Image acquisition and analysis

Z-stacks of DG images (3 frames with 2 µm interval at 20x magnification) were acquired with Nikon A1R confocal microscope (at Nikon Imaging Center, BioQuant, Heidelberg) using NIS-Elements software or with Leica SP8 confocal microscope using LAS X software. Maximum projection files of each stack were imported in Fiji (Schindelin et al., 2012), and Fos^+^ or HA-GFP^+^ neurons were manually marked after background subtraction and application of a signal threshold. Reactivation rate ((GFP^+^Fos^+^)/(GFP^+^)×100), and similarity index ((GFP^+^Fos^+^)/((GFP^+^)+(Fos^+^)-(GFP^+^Fos^+^))×100 was calculated as previously described (Cowansage et al., 2014; Gulmez Karaca et al., 2020; Milczarek et al., 2018). Data was normalized to the mean of the aged impaired group to avoid artifacts that may be caused from different viral expressions among experimental batches.

### 2.7. Statistical analysis

Blinding to experimental conditions was applied to image and behavioral analysis. Each data was subjected to a normality test (Shapiro-Wilk normality test, alpha=0.05) before further analysis. For normally distributed data ordinary one-way ANOVA followed by Tukey’s multiple comparisons test, whereas for non-normally distributed data, Kruskal-Wallis test followed by Dunn’s multiple comparisons test was used to compare three groups. When comparing two dependent data sets, paired t-test was performed if the data set showed a normal distribution, if not, Wilcoxon matched-pairs signed rank test was applied. For correlation analysis, Pearson correlation test was applied when the data followed a normal distribution. Nonparametric Spearman correlation was computed in case of a non-normally distributed dataset. Statistical analysis was performed using GraphPad prism for Mac OS X, version 8.

## 3. Results

### 3.1. Neuronal ensemble tagging in the DG of young, cognitively-impaired or -unimpaired aged mice

We aimed at characterizing DG neuronal ensembles that hold long-term representations of object location memory in order to identify possible correlates of cognitive dysfunction in the aged population. To label the neuronal ensemble activated during learning, we used the robust activity marking (RAM) system (Sørensen et al., 2016) that allowed learning-dependent expression of HA-tagged GFP (Sørensen et al., 2016) (**Figure 1A**). We have previously confirmed that this tool reliably tags the DG neuronal ensembles associated with object location memory (Gulmez Karaca et al., 2020). First, we stereotaxically delivered recombinant adeno associated viruses (rAAVs) containing the RAM-HA-GFP viral construct into the DG of young-adult (2 months old) or aged (18-20 months old) mice. Through the removal of doxycycline diet, we tagged the neuronal population activated during object location training (**Figure 1A**). When tested 24 h after training all young adult mice displayed higher than chance (50%) preference for the displaced object (**Figure 1B**), indicating intact long-term object location memory. In contrast, the performance of the aged group was heterogenous; some mice performed similarly to young mice, whereas others showed preferences closer to chance level (**Figure 1B**). To be able to characterize the aged population according to individual differences in cognitive performance, we sorted the aged group into aged impaired (AI) and aged unimpaired (AU) based on the mice’s long-term spatial memory scores (**Figure 1B**). Aged mice displaying preferences for the displaced object lower than one standard deviation below the mean of young mice (64.3%, represented with a dashed line in Figure 1B) were considered impaired. Importantly, we confirmed that all groups displayed comparable object exploration times during the training session (**Figure 1C**) and demonstrated similar locomotion and anxiety-like behavior in open-field test (**Figure 1D-F**). This indicates that poor LTM performance in AI mice was not due to lower exploratory behavior or motor or anxiety impairments.

**Figure 1.**
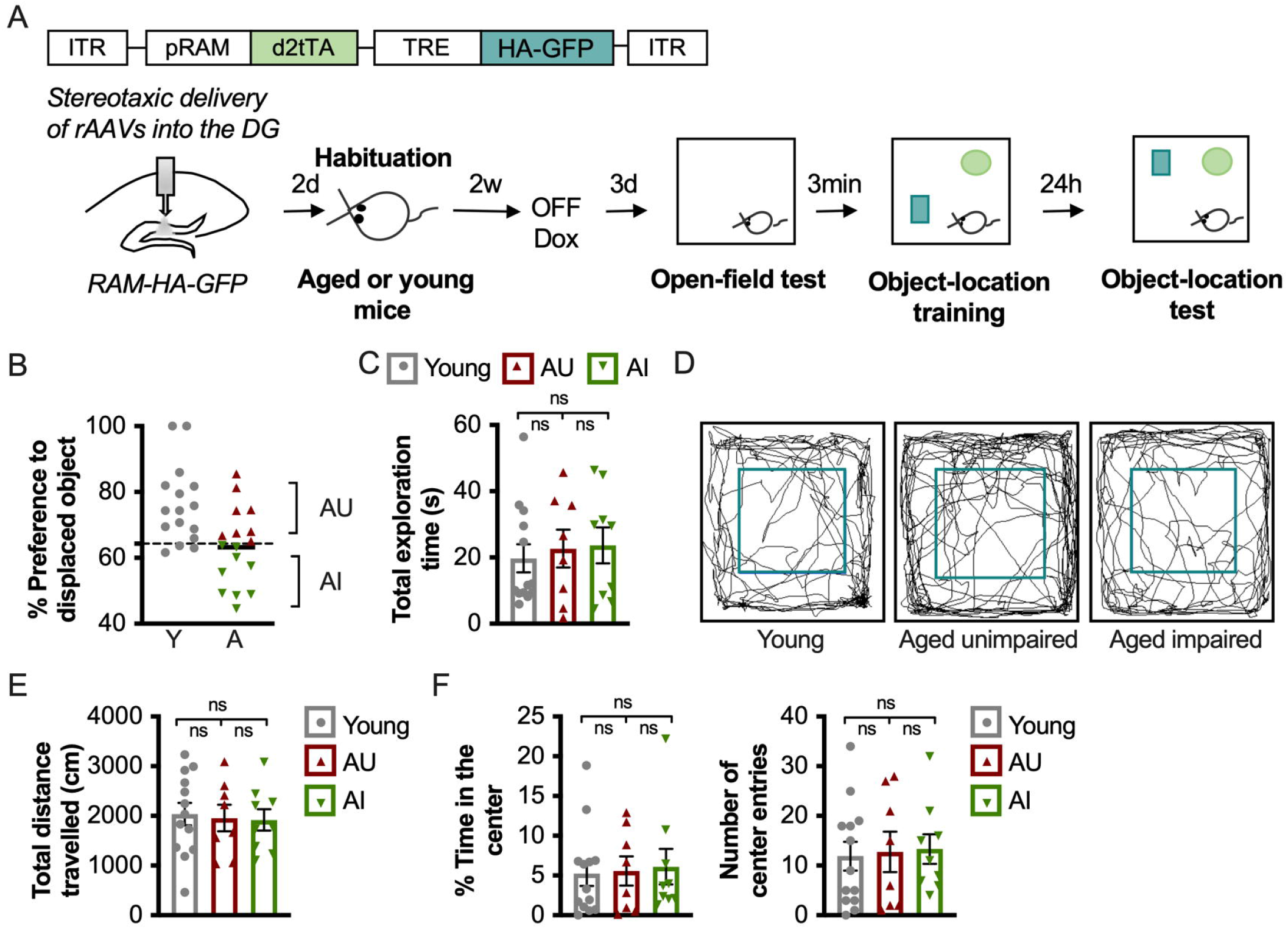
Tagging neuronal ensembles formed during spatial object-location learning in the DG of young, cognitively-impaired or -unimpaired aged mice. A) Schematic representation of the RAM-based viral construct used for neuronal ensemble tagging and of the experimental design. B) Percentage of preference for displaced object during the object location test session. Dashed line (64.3%) represents the threshold applied to categorize AI (n=9) and AU (n=8) mice (Y: mean=76.07, SD=11.80, n=16). C) Total object exploration time during the training session of object-location test (p =0.9237, Kruskal-Wallis test with Dunn’s multiple comparisons: Y (n=13) vs AU (n=8): p>0.9999; Y vs AI (n=9): p>0.9999; AU vs AI: p>0.9999). D) Representative trajectories during open-field test. E) Total distance travelled in open-field test (F(2, 27)=0.0072, p =0.9303, One-way ANOVA with Tukey’s multiple comparisons test: Y (n=13) vs AU (n=8): p=0.9694; Y vs AI (n=9): p=0.9287; AU vs AI: p=0.9936). F) Percentage of time in the center (p =0.8604, Kruskal-Wallis test with Dunn’s multiple comparisons: Y (n=13) vs AU (n=8): p>0.9999; Y vs AI (n=9): p>0.9999; AU vs AI: p>0.9999) and number of center entries (F(2,27)=0.05143, p=0.9500, one-way ANOVA with Tukey’s multiple comparisons: Y (n=13) vs AU (n=8): p=0.9826; Y vs AI (n=9): p=0.9468; AU vs AI: p=0.9926) in open-field test. ns: not significant. ITR: inverted terminal repeat, DG: Dentate gyrus of the hippocampus, Dox: doxycycline, AU: aged-unimpaired, AI: aged-impaired, Y: young, rAAV: recombinant adeno-associated viruses, TRE: tetracycline responsive element, pRAM: robust activity marking promoter. Error bars represent s.e.m.

### 3.2. Cognitively impaired aged mice exhibit impaired recall-induced neuronal ensemble reactivation in the upper blade of the DG

Next, we investigated the reactivation of spatial memory-associated neuronal ensembles in response to LTM recall. To this end, the expression of HA-GFP and Fos in the DG, that represents active neurons during learning or LTM recall respectively, was analyzed 2 h after object location test (**Figure 2A**). The neuronal population coexpressing HA-GFP and Fos represented the population active in both episodes, i.e., reactivated neuronal ensemble (**Figure 2B**). Given the previously described differences in responsiveness of the upper and lower blades of the DG to environmental exposure (Chawla et al., 2013; Gulmez Karaca et al., 2020; Marrone et al., 2012; Ramirez-Amaya et al., 2013), we analyzed the two subregions separately. We found that AI mice exhibited reduced recall-driven reactivation of the neuronal population activated by learning in the DG compared to young or AU mice (**Figure 2C**), whereas the reactivation rates observed in the DG of young and AU mice were similar (**Figure 2C**). Interestingly, the impaired reactivation rate was specific for the neuronal ensembles located in the upper, but not the lower, blade of the DG in AI mice (**Figure 2C**). Although AI mice were able to reactivate the learning-activated neuronal population at a rate significantly higher than what would occur by chance, the observed overlap above chance level was higher both in the young and AU groups compared to the AI group (**Figure 2D**). The analysis of the two DG blades independently, showed that, similarly to what we observed for the reactivation rate (**Figure 2C**), the effect originated from the upper blade (**Figure 2D**). There was no significant difference in the reactivation above chance levels of young and AU mice (**Figure 2D**). Furthermore, to evaluate whether the learning-activated neuronal ensemble pattern was reinstated equally in all of the three groups at memory recall, we applied a second formula, the similarity index (Milczarek et al., 2018). This formula complements reactivation rate by also including the number of the neurons activated solely by recall, and thereby allows to measure the degree of similarity in the neuronal activity pattern at learning and recall. We observed that AI mice exhibited the same selective impairment in the DG upper blade (**Figure 2E**). We then confirmed that the identified differences between the AI and AU were not due to differences in the number of neurons activated by learning (GFP^+^) (**Figure 2F**) or by recall (Fos^+^) (**Figure 2G**) in the aged population. Although learning triggered the activation of fewer neurons in the AU mice than the young (**Figure 2F**), this did not correlate with reactivation rate in the DG upper blade (p=0.2286, r=-0.3073, Spearman correlation, data not shown) or with memory performance (p=0.1566, r=-0.3594, Pearson correlation, data not shown) in the aged. Altogether, this set of experiments revealed that despite being able to activate a similar proportion of neurons upon learning or memory recall, cognitively impaired mice exhibit disrupted neuronal ensemble reactivation at LTM recall in the upper blade of the DG.

**Figure 2.**
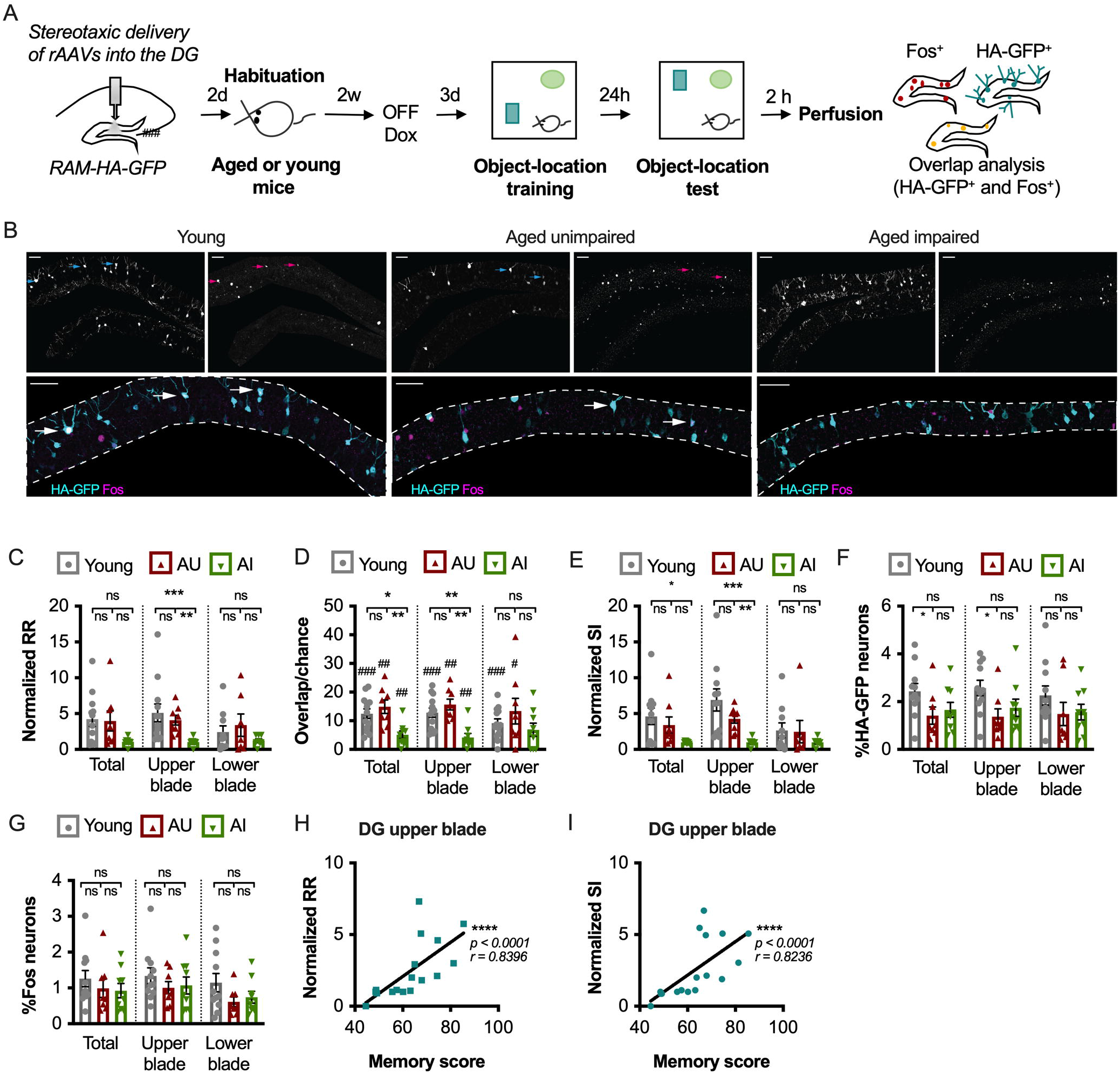
Age-related cognitive decline is associated with disrupted DG neuronal ensemble reactivation at LTM recall. A) Schematic representation of the experimental design. B) Representative images showing the immunohistochemical analysis of HA-GFP and Fos expression in young, AU and AI groups. Scale bars represent 50 µm. Dashed lines define DG granule cell layer. White arrows indicate cells with overlapping HA-GFP and Fos signal, whereas cyan and magenta arrows indicate individual HA-GFP and Fos signals, respectively. C) Normalized reactivation rates of learning-activated neuronal ensembles at memory recall (Total DG: p=0.0690, Kruskal-Wallis test with Dunn’s multiple comparisons: Y vs AU: p>0.9999; Y vs AI: p=0.1484; AU vs AI: p=0.1146; Upper blade: p=0.0003, Kruskal-Wallis test with Dunn’s multiple comparisons: Y vs AU: p>0.9999; Y vs AI: p=0.0009; AU vs AI: p=0.0016; Lower blade: p=0.8702, Kruskal-Wallis test with Dunn’s multiple comparisons: Y vs AU: p>0.9999; Y vs AI: p>0.9999; AU vs AI: p>0.9999). D) Observed-over chance-overlap between HA-GFP and Fos expressions in young, AU and AI groups (Observed-vs chance-overlap: Total Y: W = 91, p = 0.0002 by Wilcoxon test; Total AU: W = 36, p = 0.0078 by Wilcoxon test; Total AI: W = 43, p = 0.0078 by Wilcoxon test; Upper blade Y: W = 91, p = 0.0002 by Wilcoxon test; Upper blade AU: W = 36, p = 0.0078 by Wilcoxon test; Upper blade AI: W = 43, p = 0.0078 by Wilcoxon test; Lower blade Y: W = 89, p = 0.0005 by Wilcoxon test; Lower blade AU: W = 34, p = 0.0156 by Wilcoxon test; Lower blade AI: W = 33, p = 0.0547 by Wilcoxon test) (Observed/chance: Total DG: F(2,27)=7.201, p=0.0031, one-way ANOVA with Tukey’s multiple comparisons: Y vs AU: p=0.6181; Y vs AI: p=0.0152; AU vs AI: p=0.0039; Upper blade: p=0.0010, Kruskal-Wallis test with Dunn’s multiple comparisons: Y vs AU: p>0.9999; Y vs AI: p=0.0062; AU vs AI: p=0.0022; Lower blade: F(2,27)=1.336, p=0.2797, one-way ANOVA with Tukey’s multiple comparisons: Y vs AU: p=0.5108; Y vs AI: p=0.7924; AU vs AI: p=0.2557). E) Normalized similarity indices of neuronal ensemble populations activated at learning or at LTM recall (Total DG: p=0.0299, Kruskal-Wallis test with Dunn’s multiple comparisons: Y vs AU: p>0.9999; Y vs AI: p=0.0493; AU vs AI: p=0.0789; Upper blade: p=0.0002, Kruskal-Wallis test with Dunn’s multiple comparisons: Y vs AU: p>0.9999; Y vs AI: p=0.0003; AU vs AI: p=0.0050; Lower blade: p=0.6309, Kruskal-Wallis test with Dunn’s multiple comparisons: Y vs AU: p>0.9999; Y vs AI: p>0.9999; AU vs AI: p>0.9999). F) Percentage of HA-GFP^+^ neurons (Total DG: p=0.0275, Kruskal-Wallis test with Dunn’s multiple comparisons: Y vs AU: p=0.0395; Y vs AI: p=0.1667; AU vs AI: p>0.9999; Upper blade: p=0.0123, Kruskal-Wallis test with Dunn’s multiple comparisons: Y vs AU: p=0.0160; Y vs AI: p=0.1255; AU vs AI: p>0.9999; Lower blade: p=0.0842, Kruskal-Wallis test with Dunn’s multiple comparisons: Y vs AU: p=0.1364; Y vs AI: p=0.2767; AU vs AI: p>0.9999). G) Percentage of Fos^+^ neurons (Total DG: p=0.1560, Kruskal-Wallis test with Dunn’s multiple comparisons: Y vs AU: p=0.4933; Y vs AI: p=0.2275; AU vs AI: p>0.9999; Upper blade: p=0.1724, Kruskal-Wallis test with Dunn’s multiple comparisons: Y vs AU: p=0.4823; Y vs AI: p=0.2690; AU vs AI: p>0.9999; Lower blade: F(2,27)=3.460, p=0.0459, one-way ANOVA with Tukey’s multiple comparisons: Y vs AU: p=0.0665; Y vs AI: p=0.1263; AU vs AI: p=0.9298). H) Spearman correlation between the normalized neuronal ensemble reactivation in the upper blade of the DG and memory performance of aged mice at long-term object-location test (p<0.0001, r=0.8396, n=17, Spearman correlation). I) Spearman correlation between the normalized similarity index in the upper blade of the DG and memory performance of aged mice at long-term object-location test (p<0.0001, r=0.8236, n=17, Spearman correlation). In all graphs: Y (n=13), AI (n=9), AU (n=8), *p<0.05; **p<0.01; ***p<0.001; ^#^p<0.05; ^##^p<0.01; ^###^p<0.001; ns: not significant. DG: Dentate gyrus of the hippocampus, Dox: doxycycline, AU: aged-unimpaired, AI: aged-impaired, rAAV: recombinant adeno-associated viruses, pRAM: robust activity marking promoter. Error bars represent s.e.m.

### 3.3. LTM performance of aged mice correlates with the reactivation rate in the upper blade of the DG

Finally, we tested whether the neuronal ensemble reactivation rate in the upper DG blade correlates with memory performance in aged mice. For this, we performed Pearson correlation analysis between LTM performance and reactivation rate in the aged population (i.e., pooled data of AI and AU mice). Remarkably, we found a significant positive correlation between the rate of neuronal ensemble reactivation in the total DG and the memory score of the mice (p=0.0272, r=0.5396, Spearman correlation, data not shown) which primarily originated from the upper blade of the DG (**Figure 2H**), but was not present in the lower blade of the DG (p=0.3668, r=0.2496, Spearman correlation, data not shown). Similar to the reactivation rate, we observed a correlation between the degree of the similarity index in the total DG and long-term memory performance in aged mice (p=0.0064, r=0.6437, Spearman correlation, data not shown) that was selectively present in the upper blade of the DG (**Figure 2I**), but not in the lower blade of the DG (p=0.6874, r=0.1128, Spearman correlation, data not shown). Overall, these results demonstrated that cognitive performance during aging positively correlates with neuronal ensemble reactivation during the recall of LTM.

## 4. Discussion

In this study, we tagged the neuronal ensemble formed during a spatial recognition task in the DG of young, aged-impaired and aged-unimpaired mice and identified a cellular correlate of cognitive performance in aged mice. We showed that cognitively impaired aged mice exhibit impairments in recruiting the original memory engram at long-term spatial memory recall compared to young or cognitively unimpaired aged mice. We further showed that the degree of engram reactivation correlates with the strength of LTM in the aged population. We focused on long-term object location memory which has previously been shown to be impaired by aging (Oliveira et al., 2012; Wimmer et al., 2012). Increasing evidence suggests that each memory is encoded by a subset of neurons (neuronal ensembles) that are synchronously activated upon learning and reactivated by the retrieval of the memory (Josselyn et al., 2015; Tonegawa et al., 2015). A previous study by Penner and colleagues showed that neuronal reactivation upon short-term re-exposure to a previously visited context is reduced in the DG, but not CA1, of aged compared to young adult mice (Penner et al., 2011). These findings suggest that impairments in neuronal reactivation may be associated with cognitive deficits in aged individuals (Penner et al., 2011). Here, we characterized DG neuronal ensemble during the formation and retrieval of long-term recognition memory and correlated with the performance in the same task. We found that aging-related impairments in long-term object location memory are not associated with the size of the DG neuronal population activated by learning or recall, but rather with the fidelity of reactivation of the encoding population during memory retrieval. Aged impaired and unimpaired mice showed similar number of neurons activated by learning or by recall. In contrast, cognitively impaired mice had significantly disrupted reactivation rates compared to unimpaired aged, or young adult mice. Intriguingly, the differences were found in the neuronal population located in the upper, but not lower, blade of the DG. This is in line with previous studies showing that behaviorally-induced expression of activity regulated genes occurs primarily in the upper blade of the DG (Chawla et al., 2013, 2005; Erwin et al., 2020; Gulmez Karaca et al., 2020; Marrone et al., 2012; Ramirez-Amaya et al., 2013). These reports indicate that the sparse population of DG upper blade neurons form the spatial memory neuronal ensemble. Therefore, it may be expected that alterations in ensemble dynamics are specific to this subregion. Finally, our findings showed a positive correlation between the neuronal ensemble reactivation rate in the upper blade of the DG and LTM performance of aged mice. These findings suggest a novel mechanism that may underlie long-term spatial memory integrity in the aged population.

The underlying molecular and physiological causes of disrupted neuronal ensemble reactivation in aged cognitively-impaired aged mice are not understood. It is well established that aging is accompanied by anatomical and physiological changes in the DG. Namely, fewer synaptic contacts and impaired synaptic plasticity in aged versus young adults have been reported (Burke and Barnes, 2010). Interestingly, age-related impairments in LTP maintenance correlated with memory performance (Rosenzweig and Barnes, 2003). Thus, deficits in the reactivation of the original encoding neuronal population may result from impaired neuronal ensemble stabilization during memory consolidation, which could emerge from the inability to strengthen the connectivity between ensemble neurons.

Recently, a few studies showed that LTM impairments in a mouse model of Alzheimer’s disease (AD) are linked to deficits in neuronal ensemble reactivation (Perusini et al., 2017; Poll et al., 2020; Roy et al., 2016). Our findings now show that alterations in neuronal ensemble properties are a common mechanism in aging and aging-associated pathological conditions. This underscores the need for therapeutic approaches targeted at facilitating engram reactivation to restore age-associated memory impairments.

## Acknowledgements

We thank Stephanie Zeuch for critical comments on the manuscript. This work was supported by the Deutsche Forschungsgemeinschaft (DFG) [grant numbers SFB 1134 (C01), OL 437/1 and OL 437/2 to A.M.M.O.] and the Chica and Heinz Schaller foundation [fellowship and research award to A.M.M.O.]. D.V.C.B. is supported by a Landesgraduiertenförderung (LGF) completion grant (Heidelberg Graduate Academy).

## References

Brito, D.V.C., Gulmez Karaca, K., Kupke, J., Mudlaff, F., Zeuch, B., Gomes, R., Lopes, L. V., Oliveira, A.M.M., 2020a. Modeling human age-associated increase in Gadd45γ expression leads to spatial recognition memory impairments in young adult mice. Neurobiol. Aging. https://doi.org/10.1016/j.neurobiolaging.2020.06.021

Brito, D.V.C., Kupke, J., Karaca, K.G., Zeuch, B., Oliveira, A.M.M., 2020b. Mimicking age-associated Gadd45γ dysregulation results in memory impairments in young adult mice. J. Neurosci. 40, 1197–1210. https://doi.org/10.1523/JNEUROSCI.1621-19.2019

Burke, S.N., Barnes, C.A., 2010. Senescent synapses and hippocampal circuit dynamics. Trends Neurosci. https://doi.org/10.1016/j.tins.2009.12.003

Burke, S.N., Barnes, C.A., 2006. Neural plasticity in the ageing brain. Nat. Rev. Neurosci. https://doi.org/10.1038/nrn1809

Chawla, M.K., Guzowski, J.F., Ramirez-Amaya, V., Lipa, P., Hoffman, K.L., Marriott, L.K., Worley, P.F., McNaughton, B.L., Barnes, C.A., 2005. Sparse, environmentally selective expression of Arc RNA in the upper blade of the rodent fascia dentata by brief spatial experience. Hippocampus 15, 579–586. https://doi.org/10.1002/hipo.20091

Chawla, M.K., Penner, M.R., Olson, K.M., Sutherland, V.L., Mittelman-Smith, M.A., Barnes, C.A., 2013. Spatial behavior and seizure-induced changes in c-fos mRNA expression in young and old rats. Neurobiol. Aging 34, 1184–1198. https://doi.org/10.1016/j.neurobiolaging.2012.10.017

Cowansage, K.K., Shuman, T., Dillingham, B.C., Chang, A., Golshani, P., Mayford, M., 2014. Direct Reactivation of a Coherent Neocortical Memory of Context. Neuron. https://doi.org/10.1016/j.neuron.2014.09.022

Erwin, S.R., Sun, W., Copeland, M., Lindo, S., Spruston, N., Cembrowski, M.S., 2020. A Sparse, Spatially Biased Subtype of Mature Granule Cell Dominates Recruitment in Hippocampal- Associated Behaviors. Cell Rep. 31. https://doi.org/10.1016/j.celrep.2020.107551

Gulmez Karaca, K., Brito, D.V.C., Zeuch, B., Oliveira, A.M.M., 2018. Adult hippocampal MeCP2 preserves the genomic responsiveness to learning required for long-term memory formation. Neurobiol. Learn. Mem. 149, 84–97. https://doi.org/10.1016/j.nlm.2018.02.010

Gulmez Karaca, K., Kupke, J., Brito, D.V.C., Zeuch, B., Thome, C., Weichenhan, D., Lutsik, P., Plass, C., Oliveira, A.M.M., 2020. Neuronal ensemble-specific DNA methylation strengthens engram stability. Nat. Commun. 11. https://doi.org/10.1038/s41467-020-14498-4

Ianov, L., Rani, A., Beas, B.S., Kumar, A., Foster, T.C., 2016. Transcription profile of aging and cognition-related genes in the medial prefrontal cortex. Front. Aging Neurosci. 8. https://doi.org/10.3389/fnagi.2016.00113

Josselyn, S.A., Köhler, S., Frankland, P.W., 2015. Finding the engram. Nat. Rev. Neurosci. https://doi.org/10.1038/nrn4000

Marrone, D.F., Satvat, E., Shaner, M.J., Worley, P.F., Barnes, C.A., 2012. Attenuated long-term Arc expression in the aged fascia dentata. Neurobiol. Aging 33, 979–990. https://doi.org/10.1016/j.neurobiolaging.2010.07.022

Milczarek, M.M., Vann, S.D., Sengpiel, F., 2018. Spatial Memory Engram in the Mouse Retrosplenial Cortex. Curr. Biol. https://doi.org/10.1016/j.cub.2018.05.002

Oliveira, A.M.M., Hemstedt, T.J., Bading, H., 2012. Rescue of aging-associated decline in Dnmt3a2 expression restores cognitive abilities. Nat. Neurosci. 15, 1111–1113. https://doi.org/10.1038/nn.3151

Penner, M.R., Roth, T.L., Chawla, M.K., Hoang, L.T., Roth, E.D., Lubin, F.D., Sweatt, J.D., Worley, P.F., Barnes, C.A., 2011. Age-related changes in Arc transcription and DNA methylation within the hippocampus. Neurobiol. Aging. https://doi.org/10.1016/j.neurobiolaging.2010.01.009

Perusini, J.N., Cajigas, S.A., Cohensedgh, O., Lim, S.C., Pavlova, I.P., Donaldson, Z.R., Denny, C.A., 2017. Optogenetic stimulation of dentate gyrus engrams restores memory in Alzheimer’s disease mice. Hippocampus. https://doi.org/10.1002/hipo.22756

Poll, S., Mittag, M., Musacchio, F., Justus, L.C., Giovannetti, E.A., Steffen, J., Wagner, J., Zohren, L., Schoch, S., Schmidt, B., Jackson, W.S., Ehninger, D., Fuhrmann, M., 2020. Memory trace interference impairs recall in a mouse model of Alzheimer’s disease. Nat. Neurosci. https://doi.org/10.1038/s41593-020-0652-4

Ramirez-Amaya, V., Angulo-Perkins, A., Chawla, M.K., Barnes, C.A., Rosi, S., 2013. Sustained transcription of the immediate early gene arc in the dentate gyrus after spatial exploration. J. Neurosci. 33, 1631–1639. https://doi.org/10.1523/JNEUROSCI.2916-12.2013

Rosenzweig, E.S., Barnes, C.A., 2003. Impact of aging on hippocampal function: Plasticity, network dynamics, and cognition. Prog. Neurobiol. https://doi.org/10.1016/S0301-0082(02)00126-0

Roy, D.S., Arons, A., Mitchell, T.I., Pignatelli, M., Ryan, T.J., Tonegawa, S., 2016. Memory retrieval by activating engram cells in mouse models of early Alzheimer’s disease. Nature. https://doi.org/10.1038/nature17172

Schindelin, J., Arganda-Carreras, I., Frise, E., Kaynig, V., Longair, M., Pietzsch, T., Preibisch, S., Rueden, C., Saalfeld, S., Schmid, B., Tinevez, J.Y., White, D.J., Hartenstein, V., Eliceiri, K., Tomancak, P., Cardona, A., 2012. Fiji: An open-source platform for biological-image analysis. Nat. Methods. https://doi.org/10.1038/nmeth.2019

Small, S.A., Chawla, M.K., Buonocore, M., Rapp, P.R., Barnes, C.A., 2004. Imaging correlates of brain function in monkeys and rats isolates a hippocampal subregion differentially vulnerable to aging. Proc. Natl. Acad. Sci. U. S. A. https://doi.org/10.1073/pnas.0400285101

Small, S.A., Schobel, S.A., Buxton, R.B., Witter, M.P., Barnes, C.A., 2011. A pathophysiological framework of hippocampal dysfunction in ageing and disease. Nat. Rev. Neurosci. https://doi.org/10.1038/nrn3085

Sørensen, A.T., Cooper, Y.A., Baratta, M. V., Weng, F.J., Zhang, Y., Ramamoorthi, K., Fropf, R., Laverriere, E., Xue, J., Young, A., Schneider, C., Gøtzsche, C.R., Hemberg, M., Yin, J.C.P., Maier, S.F., Lin, Y., 2016. A robust activity marking system for exploring active neuronal ensembles. Elife 5. https://doi.org/10.7554/eLife.13918

Tonegawa, S., Pignatelli, M., Roy, D.S., Ryan, T.J., 2015. Memory engram storage and retrieval. Curr. Opin. Neurobiol. https://doi.org/10.1016/j.conb.2015.07.009

Wimmer, M.E., Hernandez, P.J., Blackwell, J., Abel, T., 2012. Aging impairs hippocampus-dependent long-term memory for object location in mice. Neurobiol. Aging. https://doi.org/10.1016/j.neurobiolaging.2011.07.007

Zhang, S.J., Steijaert, M.N., Lau, D., Schütz, G., Delucinge-Vivier, C., Descombes, P., Bading, H., 2007. Decoding NMDA Receptor Signaling: Identification of Genomic Programs Specifying Neuronal Survival and Death. Neuron. https://doi.org/10.1016/j.neuron.2007.01.025

